# Designing minimal *E. coli* genomes using variational autoencoders

**DOI:** 10.1101/2024.10.22.619620

**Authors:** Anastasiia Shcherbakova, Kieren Sharma, Yige Chen, Claire S. Grierson, Lucia Marucci, Daniel Buchan, Chris P. Barnes

## Abstract

Designing minimal bacterial genomes remains a key challenge in synthetic biology. There is currently a lack of efficient tools for the rapid generation of streamlined bacterial genomes, limiting research in this area. Here, using a pangenome dataset for *Escherichia coli*, we show that variational autoencoders with modified loss functions can successfully create minimised genomes retaining the essential genes identified in the literature. We then sampled new genomes from our fitted model and performed computational validation using an *E. coli* whole-cell model. We found 6 out of 100 of the sampled genomes were viable in the computer model. These underwent a minimization routine starting from the MG1655 genome giving rise to six new minimal genomes with around a 40 % reduction in size. This study proposes a rapid, machine learning-based approach for bacterial sequence generation, that could accelerate the genomic design process.

## 1 Introduction

Research on reduced genomes is important for understanding biological principles, revealing the minimal set of genes essential for cell survival and for industrial purposes [1]. A number of interesting outcomes have arisen from the creation of bacteria with reduced genomes including increased growth rate and biomass production [2, 3, 4, 5], increased gene product output [6, 7, 8] and enhanced cellular stability[9]. In addition to the increase in growth rate and decrease in byproduct production, Mogger-Reischer *et. al*. have also reported that cells engineered with only essential genes show faster evolution in terms of relative fitness compared to wild-type *Mycoplasma mycoides* cells [10]. This finding may suggest that genome reduction may also enhance bacteria’s adaptability and evolutionary potential, further supporting its utilisation in biotechnological and research settings.

There are two primary approaches to creating new minimal genomes: The bottom-up approach involves *de novo* synthesis of a new artificial genome allowing for the complete redesign of bacterial genomes from scratch [11]. The top-down approach involves the reduction of existing wild-type genomes by deleting non-essential genetic elements. Despite the bottom-up approach being a success, the bacterial design principles remain unknown, posing challenges for completely new assemblies.

Several top-down reduced genome strains have been developed from the *Escherichia coli* strains K-12 W3110 and K-12 MG1655, each tailored for specific biotechnological applications [12]. One such strain, the Minimal Genome Factory (MGF-01), was designed to contain only the essential genes necessary for fermentative production. MGF-01 lacks 22% of the original K-12 W3110 genome while remaining comparable to the wild-type strain in many aspects [13]. Notably, MGF-01 accumulated higher L-threonine levels than the wild-type strain, supporting the hypothesis that genome reduction can lead to increased productivity [14]. Another key development is the Multiple Deletion Series (MDS), designed to focus on genetic stability and metabolic efficiency to optimise the insertion of foreign genes for industrial purposes [15]. The MDS series was derived from the MG1655 strain by eliminating insertion sequences and reducing additional genetic components to create strains such as MDS12, MDS41, MDS43, and MDS69 [12]. Of these, MDS41 and MDS42 exhibited no significant differences in growth rates compared to the wild-type strain [16], although MDS69 showed a nearly 20% reduction in growth rate relative to MG1655 [17]. Despite its slower growth, MDS42 demonstrated a significant increase 83% more L-threonine than the engineered MG1655, highlighting that genome reduction can enhance the production of specific compounds even when growth is affected [8]. The smallest *E. coli* genome reported so far is Δ33a which has a size of 2.83Mb [18]. This reduced strain showed sensitivity to oxidative stress but subsequent work showed this could be alleviated using adaptive laboratory evolution [19].

Increasingly, computational modelling and machine learning techniques have been used for minimal genome design including the use of whole cell models [20, 21, 22]. Recently variational autoencoders (VAEs), generative models combining deep neural networks and probabilistic modelling [23], were used for capturing genomic variation. These were used to simulate human genomes [24, 25] and for exploration and design in bacterial genome space [26]. In this work we focus on developing a genome variation model for *E. coli* using a collection of high quality genomes comprising a large-scale pangenome [27]. We use the presence-absence gene matrix to train a VAE and explore different loss functions to capture variation such that the resultant model produces reduced genomes (Figure 1). Finally, we develop a greedy algorithm to move from simulated binary genomes to actual reduced genomes. We show that our generated genomes have a size comparable and smaller that the state of the art in experimental minimal *E. coli* genomes. This approach also provides a means for a systematic experimental programme to achieve reduced and minimal *E. coli* genomes using data-driven generative models.

**Figure 1.**
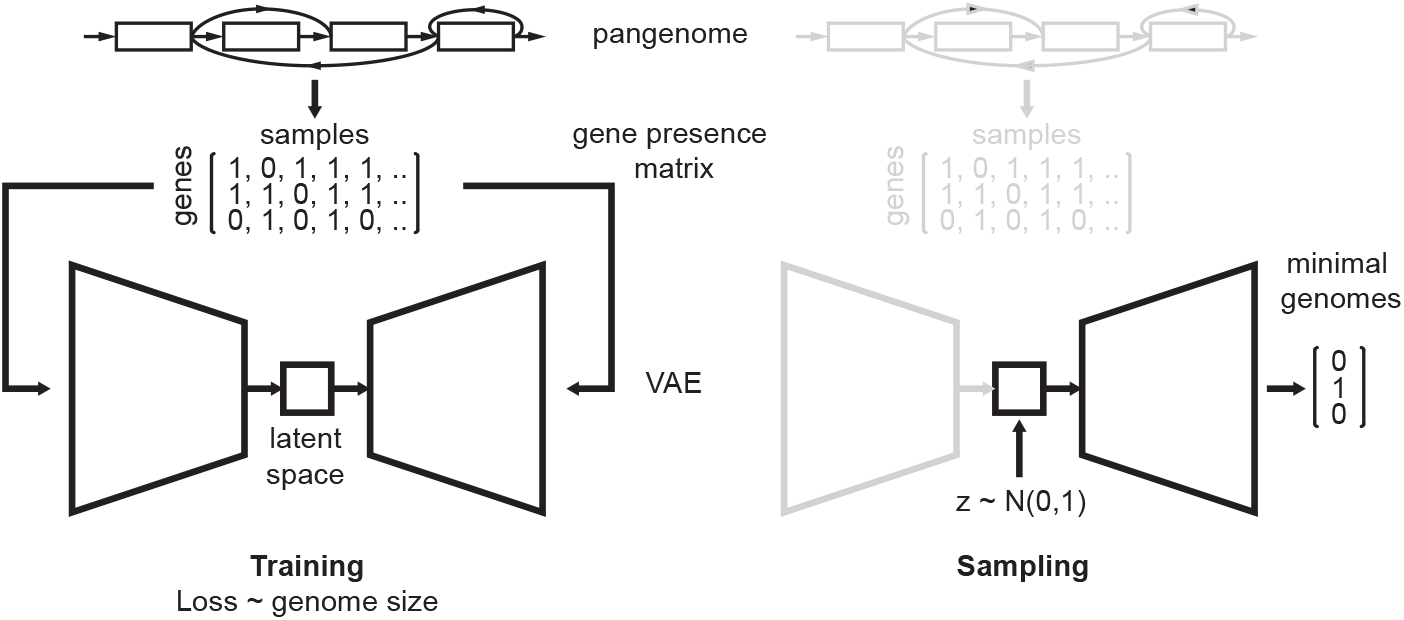
Overview of the approach to minimal genome generation. In the training stage, a variational autoencoder is fit to the binary gene presence matrix with a loss term proportional to genome size. In the generation or sampling stage, new genomes are sampled from the fitted model.

## 2 Results

### 2.1 Data Properties

The data set comprises a curated collection of *E. coli* and *Shigella* genomes [27]. We used as input the presence-absence binary matrix. The matrix consists of rows representing the gene names (55,039 total), columns representing the sample names (7,511 total) and one column representing the sample phylogroup. The data was pre-processed by removing the samples with no metadata provided, reducing the samples from 7,511 to 5,953. Figure 2a shows the distribution of genome size in terms of gene number. The maximum and minimum genome sizes are approximately 3800 and 5,300 genes respectively. Figure 2b visualises the gene frequency and, as expected, there is a collection of genes that are present across most of the genomes, which indicates the core genome. There were also many genes that occurred with very low frequency, indicating heterogeneous accessory genomes across isolates. Although, previous analyses have removed genes with low frequencies [26], we decided to keep all the genes, even the low frequency ones, for training the models. We also checked the abundance of 358 essential genes in the *E. coli* genome as reported by Goodall *et. al*. [28] over all genomes. In the Horesh *et. al*. dataset, some genes were split into multiple gene fragments. We aggregated these fragments into a single representation for each gene and the dataset contained only 328 essential genes, with 30 not present in the data set. The essential gene abundance distribution is given in Figure 2c. Most genomes contain a high number of essential genes; however, interestingly, a much smaller portion does not have all the essential genes, with the minimal number being 284. This possibly reflects both the ambiguous concept of essential genes, which depends on the environment and genetic background [28], and also the fact that many of these strains are members of a complex mammalian gut microbiome. Figure 2d shows the first two principal components of the data, coloured by phylogroup. PC1 separates phylogroup E and the rest of the phylogroups. PC2 separates the B1 group from the E group. As described by Horesh *et. al*. [27], all phylogroup E lineages contain *Enterohemorrhagic Escherichia coli (EHEC)* isolates.

**Figure 2.**
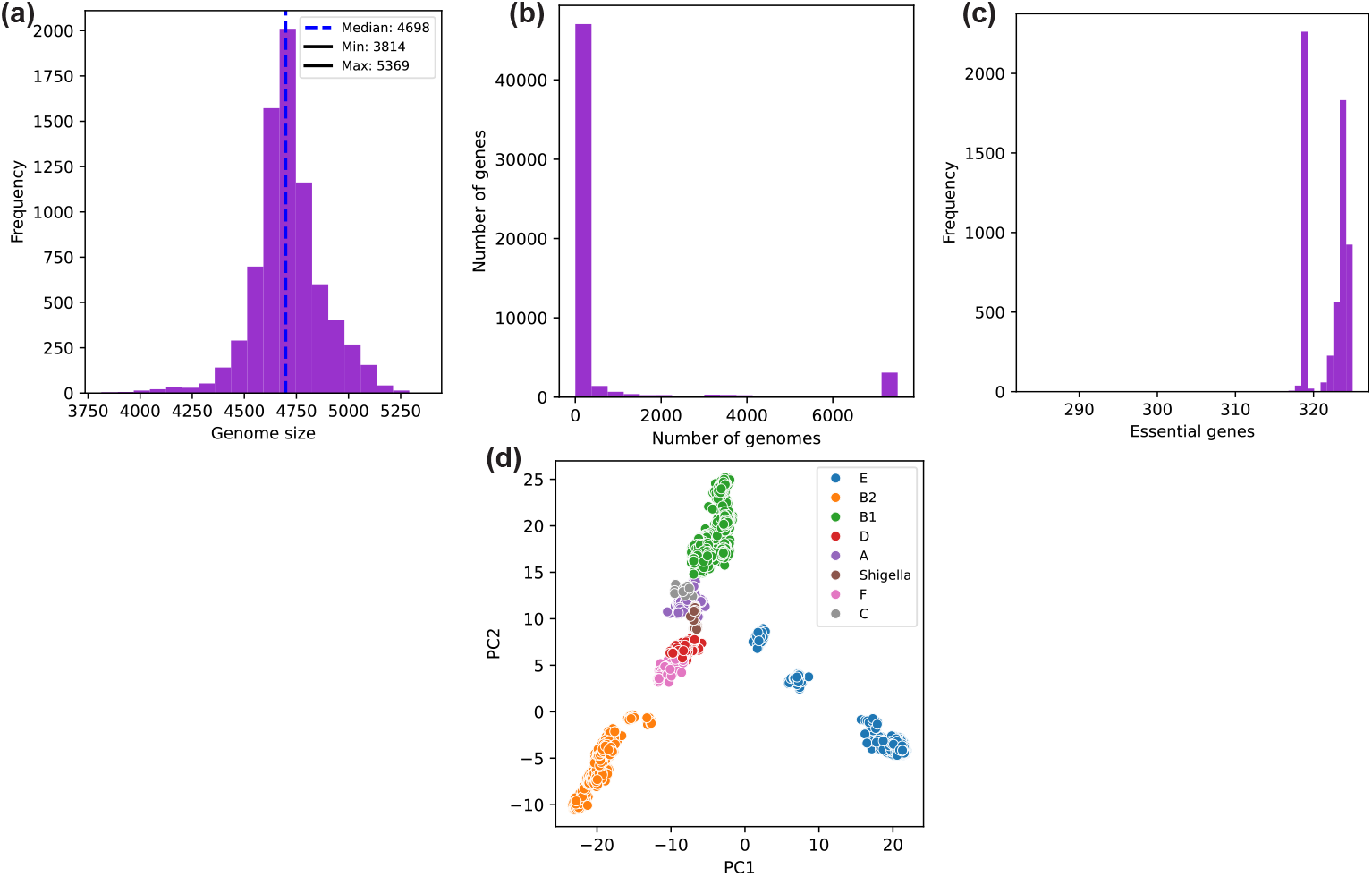
Properties of the genomes in the *E. coli* pangenome dataset. (a) Distribution of genome size. (b) Number of genes vs number of genomes, representing the frequency of genes across the genomes. (c) Frequency of essential genes. (d) PCA of the dataset coloured by the phylogroup.

### 2.2 Exploring loss functions to capture minimal genomes

We developed a number of models that differed in how the genome variation was captured (see Methods). The VAE model’s the posterior probability distribution, *p*(*z*|*x*) where *x* is the data (set of all genomes) and *z* is the latent space, a compressed representation of the genome information. VAEs are fit using variational inference, which is a type of approximate Bayesian inference where the posterior distribution is approximated by a family of tractable distributions (here we used multivariate Gaussian distributions). The model balances how well the model can generate genomes (the reconstruction loss) with how close the model resembles the prior distribution (the KL divergence loss).

Our first model (v0) used a standard *β−*VAE loss function with terms for reconstruction loss and KL divergence. The *β* refers to a weighting of the relative importance of these two loss terms. In version v1 we added another loss term that was proportional to the total gene number. By adding this term, the model tries to capture the genome variation using fewer genes, thus capturing the most common genes across the dataset (i.e. the core genome) and ignoring the less common genes (i.e. the accessory genome). A challenge with fitting these models is that they can become stuck in local minima, so how the fitting is performed is important. There are various ways to move from simpler models to the final more complex model through changing the relative weighting of the terms in the loss function (known as annealing). In version v2 we explored different ways to anneal, in particular we used a cyclical annealing for the beta coefficient which modified the KL divergence influence. The results are shown in Figures S1-S4.

Comparing the base model v0 with v1, we can notice more clustered points on the latent space by phylogroup and reduction in sparsity. The genome size distribution considerably changed in size as intended, with the median reducing from around 4500 to 3950 genes in a genome. The number of essential genes stayed relatively similar with the addition of the total genes term, with a slight reduction in minimum and maximum values. F1 score distribution remained the same with the loss function change. This shows that the model reconstruction ability was not compromised despite adding a total gene number term. Comparing v1 to v2, there is no considerable difference in the genome size, essential gene number or F1 score distribution. However, a significant alteration occurred in the model’s latent space. The clustered latent space in v1 became more sparse and there is a clear vertical division of phylogroups B2, B1 and E. An increase in sparsity and a decrease in the mixing of the points from different phylogroups allowed for the production of more meaningful latent space, prompting us to use the method for the final model.

We next explored further modifying the influence of the total gene number term. We experimented with both static weights and gamma coefficient scheduling to achieve these results (see Methods, Equation 3). We found that using static weights above 2.5 caused the model to produce all-zero outputs, likely because the corresponding loss term became too dominant. However, when we used gamma scheduling, transitioning from 2.0 to 0.1 instead of from 1.0 to 0.1, we achieved successful training. The best combination we found is *ω* = 1, *γ* : 2 *→* 1, denoted model v3. The results for this model are presented in Figure 3 and Figure S5. The sampled genomes from the final model show a considerable reduction in genome size, with a median distribution of around 3100 genes per genome (Figure 3a). The latent space provides good separation between phylogroups while the genomes retain a good median number of essential genes (Figure 3b, c respectively). In addition, within the distribution of genomes, there is good retention of essential genes even for genomes down to sizes of around 2500 genes (Figure 3d).

**Figure 3.**
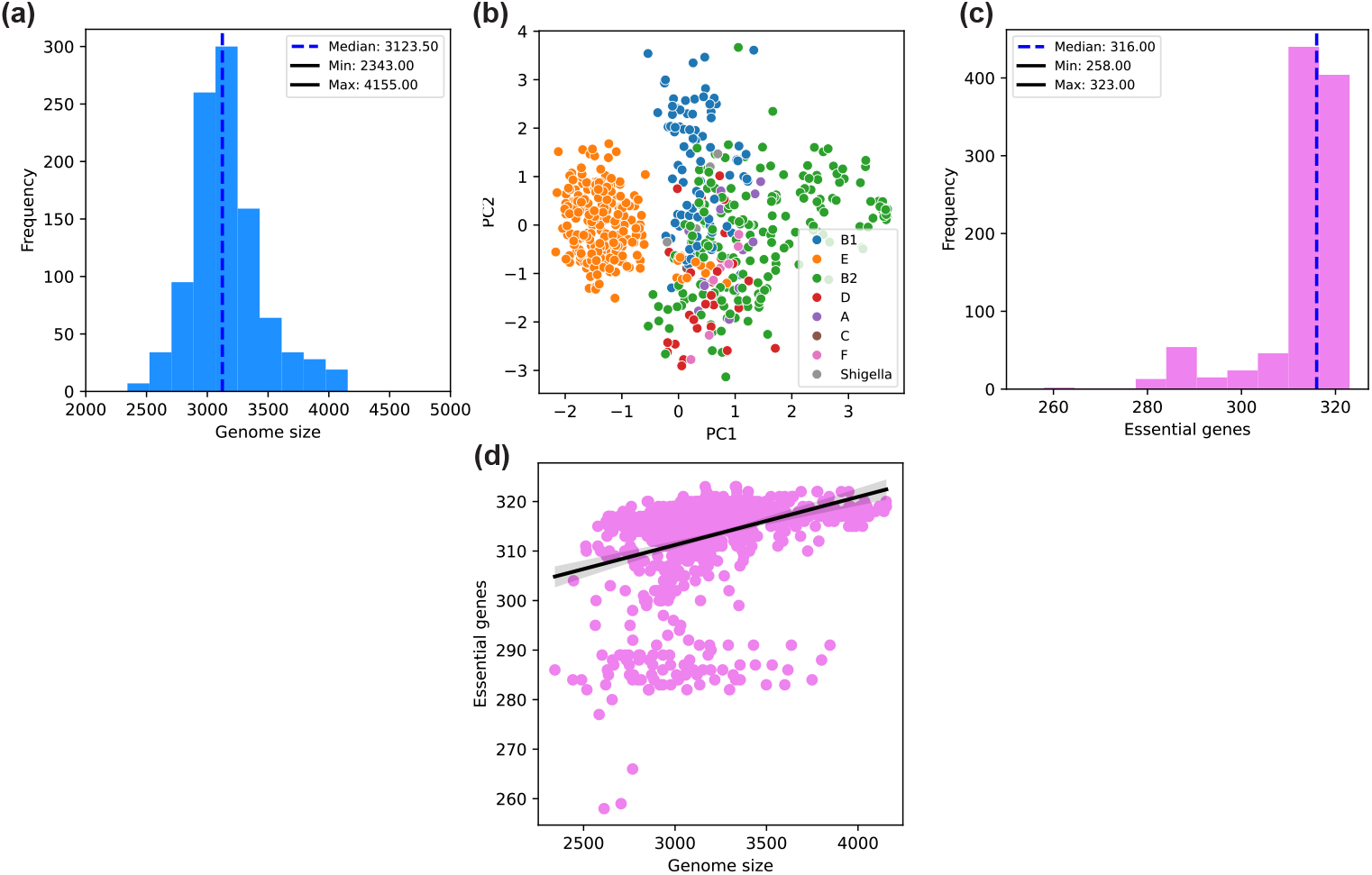
Properties of 1000 sampled genomes from the final model (v3). (a) Distribution of genome size. (b) Latent space visualisation. (c) Distribution of the number of essential genes. (d) Genome size vs number of essential genes.

### 2.3 Validation of sampled genomes using an *E. coli* whole-cell model

The *E. coli* whole-cell model (WCM) is a computer simulation that integrates many cellular processes—such as metabolism, gene regulation, and protein activity—into a single framework to predict how the bacterium functions as a whole [29, 30]. WCMs have previously been used in the context of minimal genome design as a computational validation step [20, 21, 22]. The *E. coli* WCM integrates primary data from three laboratory strains (K-12 MG1655, B/r, and BW25113) and therefore models a “hybrid” genome consisting of 1,872 genes. To first assess the model’s ability to capture experimental gene essentiality, we simulated a genome-wide single-gene knockout screen and compared the resulting *in silico* essentiality classifications to experimental CRISPR–based fitness measurements from Rousset *et al*. [31], yielding 90.2% accuracy (95.3% specificity, 51.6% sensitivity). This provides us with experimental grounding for using the WCM as a computational screening tool for minimal genome designs, while recognising that it does not validate synergistic effects arising from higher-order combinatorial deletions.

To validate our simulated genomes using the WCM, we first assessed the *in silico* viability of 50 representative genomes, one per lineage among the 50 largest PopPUNK lineages in Horesh *et al*.. Since the version of the WCM we used mechanistically models only approximately 73% of the well-characterised MG1655 genes [32], we first restricted each binary presence–absence genome vector to this set of genes that we term the *WCM gene set*. In addition, before simulation we restored any missing WCM-essential genes—identified using an *in silico* genome-wide single-gene deletion screen, because, by definition, absence of any such gene renders the model cell non-viable. Among lineage genomes, the number of restored WCM-essential genes ranged from 8–13 (mean 8.43). All 50 lineage genomes were viable, each completing 20 generations, where one generation denotes a complete celldivision cycle.

We next validated 100 genomes sampled from the v3 model’s latent space. We followed the same procedure as above, restricting to the WCM gene set and adding back missing WCM-essential genes prior to simulation. After filtering, the minimal genomes retained 63-76% of genes, and the number of essential genes restored per sampled genome ranged from 15–48 (mean 21.4). Under the same viability criterion (20 generations), six genomes were viable. Among the 94 non-viable designs, the mean generations completed was 3.2. To confirm robustness, we re-simulated the six viable designs with three independent random seeds to initialise stochastic processes in the WCM. Four of six remained viable in all repeats, while two designs (samples 19 and 40) were viable in 2/3 repeats and failed in 1/3, and are therefore considered repeat-inconsistent.

Phenotypic analysis of the six viable minimal-genome designs against 10 wild-type (WT) repeats was then performed (Figure 4). Figure 4a shows the size of the genomes restricted to the WCM gene set. Figure 4b shows the specific growth rate vs time, of the simulations where trajectories largely overlapped the WT envelope, with one design (Sample 40) exhibiting a sustained lower trajectory. Similarly, when comparing cell mass with WT (Figure 4c), the minimal genomes were again congruent with WT except Sample 40, which tracked below the cohort. Comparing mean cell mass vs mean specific growth rate across all 100 sampled designs relative to WT means (Figure 4c), a positive correlation was observed. The four repeat-stable designs clustered closely to the WT point, while genome 40 lay at the lower-mass/lower-growth side of WT (mean specific growth rate 0.468 *h*^*−*1^; mean cell mass 1158.28 fg). We also explored the growth rate as a function of genome size, which showed a positive correlation (Figure 4e) and the relationship between the viability and the size of the original sampled genomes (Figure 4f). Together, these results indicate that the repeat-stable WCM-viable minimal designs sampled by the VAE can match WT physiological trajectories (growth and biomass) despite substantial gene set reductions, consistent with generating minimal genomes while preserving core essential functions.

**Figure 4.**
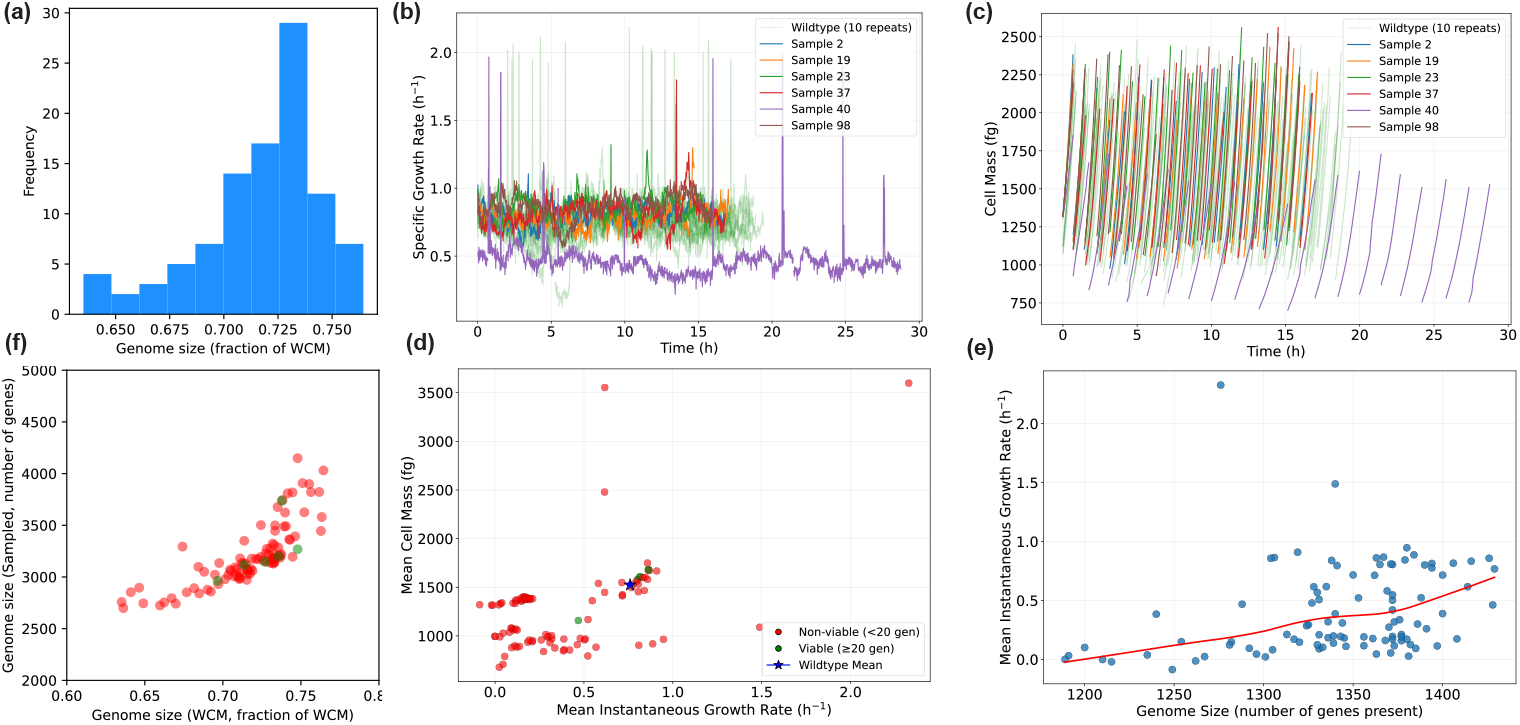
*E. coli* whole cell model validation. (a) Size of the sampled genomes restricted to the WCM gene set. b) Specific growth rate of viable genomes compared to the wildtype. (c) Cell mass of viable genomes compared to the wildtype. (d) Specific growth rate versus cell mass. (d) Specific growth rate as a function of genome size. (f) Size of the WCM genomes vs size of the sampled genomes.

### 2.4 Realizing minimal *E. coli* genome sequences

We took the six sampled genomes that were successfully validated in the WCM and explored their overlap and diversity. Figure 5a and b show the pairwise gene distance matrix and the resultant phylogenetic tree respectively. There is limited overlap in the genomes with differences ranging from around 630 to 1200 genes. To move from gene space into sequence space, we developed a greedy minimisation algorithm that iteratively removes gene sequences starting from an original genome (see Methods). We applied this algorithm to the six validated genomes starting from a wildtype MG1655 genome. Figure 5c shows the resultant size of the genomes and the percentage reduction from the wildtype. The genomes range from 2,686,277 bp to 2,867,946 bp or 42.1% to 38.2% reduction.

**Figure 5.**
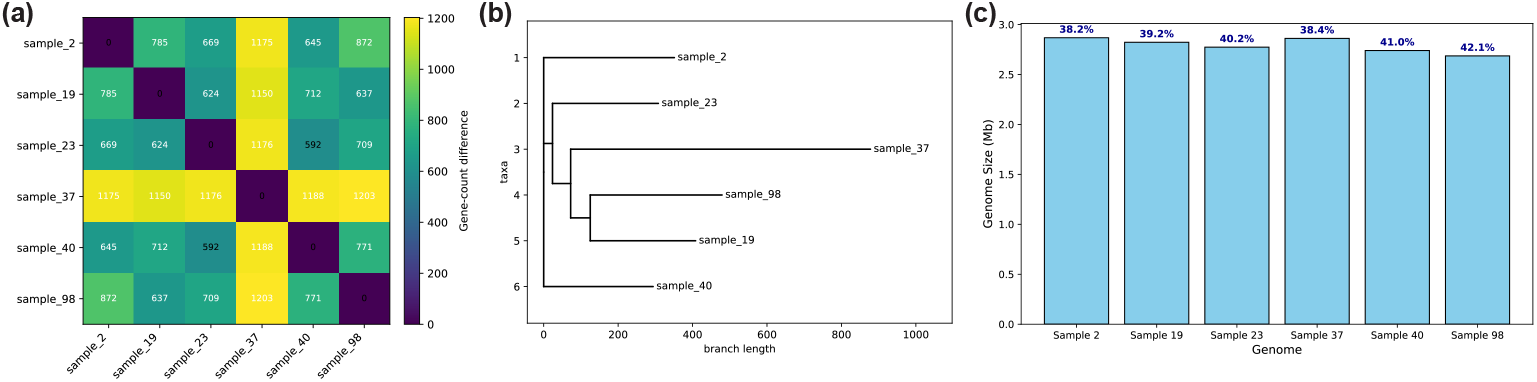
Realizing six minimal *E. coli* genome sequences. (a) Distance matrix between the six final genomes. (b) Phylogenetic tree showing the distance relationship. (c) Size distribution in Mb of the final genomes, processed through the reduction of the MG1655 *E. coli* genome.

## 3 Discussion

We demonstrated the ability of a VAE to generate new minimised bacterial *E. coli* genomes. We first fit a VAE model to around 6000 *E. coli* genomes comprising a pangenome gene presence-absence dataset. The VAE model included a modified loss term proportional to genome size. By sampling latent vectors from the fitted model and passing through the decoder part of the network, we were able to generate sets of genes corresponding to minimized genomes. We then took 100 sampled genomes and validated them in an *E. coli* WCM. Six out of the hundred genomes were viable. These viable genomes were then converted into actual sequence using a greedy minimization approach. The genomes ranged from 2.6Mb to 2.8 Mb, comprising a reduction of 40% which is consistent with the current smallest minimised E. coli genome of 2.8 Mb [18]. These results highlight the potential of this approach to create more efficient and streamlined genomes for industrial bioprocessing, where smaller genomes can lead to reduced metabolic burden and increased production yields. Our work extends the efforts of Dudek and Precup [26], who also explored machine learning driven bacterial genome generation using VAEs. While they focused on generating genomes across various bacterial species and faced challenges with small and unbalanced datasets, we concentrated solely on *E. coli* to address specific industrial and research applications. To achieve that, we used a large high quality dataset [27]

The outcome of the minimal genome generation experiments also shows a path to new genome reduction algorithms. A key challenge in genome minimisation is determining the organism’s viability after gene deletions. In the future, by building a phylogenetic tree of the generated genomes, we can in principle estimate the genome at internal nodes of the tree. There is therefore potential to traverse the tree in a greedy fashion, selecting the internal node with the largest number of descendants, removing those genes and then verifying the viability of the genome experimentally. If the genome is viable, we continue to the next internal node below; if that genome is not viable we move to the adjacent node with the next largest number of descendants. Future work will focus on validating this hypothesis and developing computational and experimental tools to implement this strategy.

There are a number of limitations. We used a WCM gauge whether the genomes were realistic, but ultimately, the generated genomes will need to be experimentally verified, which is currently a time consuming and costly process [1]. New experimental approaches for engineering the *E. coli* genome have been developed [33], and our modelling approach could help guide future attempts. We used a large data set of *E. coli* genomes which provides limited genetic diversity. In the future, data could be combined from multiple closely related species in order to increase genetic diversity. Another limitation of our approach was the limited hyperparameter tuning. Although we explored hidden dimensions, latent dimensions, and the learning rate, in the future, a thorough exploration using Bayesian optimisation should be performed. In conclusion, this study has demonstrated the potential of using generative models to design bacterial genomes and provides foundations for AI approaches to engineering biology.

## 4 Methods

### 4.1 VAE model

#### 4.1.1 Basic VAE model (v0)

All the VAE models were implemented using the PyTorch framework. The encoder consists of three fully connected layers with a ReLU activation function and additional batch normalisation. The first layer contains the input dimension of the total number of features (genes number) and a hidden dimension of 1,024 for the basic model and 512 for the rest. The output of the encoder is the mean and log variance for the latent space. The VAE also employs a reparametrisation trick that allows sampling to be differentiable.

The decoder consists of four fully connected layers. The first 3 layers contain the ReLU activation function and batch normalisation. The last layer contains the Sigmoid activation function, which gives outputs between 0 and 1. The first layer includes the same input and output dimensions as the encoder. To make the training process more efficient, the model uses Xavier initialisation for the weights of the linear layers and initialised biases to zero. The model diagram can be seen in Figure S2. The loss function of the base VAE consisted of the sum of reconstruction loss and KL divergence loss and is given by

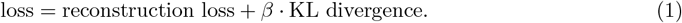

The KL term is annealed linearly through a beta term from 0.1 to 1.0 (3.2). Gradient clipping was used for all VAE model training to prevent the problem of exploding gradients, which occurs when gradients become large during backpropagation. This problem is common in deep architecture with exponentially growing gradients. The gradients were clipped to a maximum value of 1.0, which ensured stable model training.

#### 4.1.2 VAE models with modified loss function (v1)

The first modification of the loss function included a term for the total gene number, given by

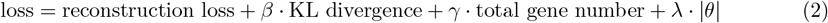

where gamma is linearly annealed from 1.0 to 0.1. L1 regularisation was used to prevent overfitting and improve generalisation. The lambda coefficient to control the regularisation was chosen to be 0.01.

#### 4.1.3 VAE model with cyclic annealing (v2)

Previously, the KL divergence term followed a linear annealing schedule. Linear annealing is not always the optimal schedule because it can apply too much regularisation too early. Early regularisation hinders models’ ability to explore the latent space and leads to poor reconstruction [34]. Therefore, the KL divergence term was annealed to cyclic annealing for more optimal learning of the latent space. Such an annealing schedule introduces high and low regularisation periods, letting the model explore different regions of the latent space. In this case, the annealing follows the cosine structure, where the weight of the KL divergence term changes periodically following the cosine function, ramping up and down. The cyclic annealing schedule aims to enhance the learning of the latent space, ensuring a balanced trade-off between reconstruction accuracy and latent space regularisation. The loss function for this model is given by Equation 2 and included L1 regularisation.

#### 4.1.4 VAE model with modified loss function, cyclic annealing and additional weights (v3)

The final model includes all the modifications from VAE v2. However, it also includes an additional weight *ω* for the total gene number term to see how the increase in the influence of that term reduced the genome size distribution. The final loss function was given by

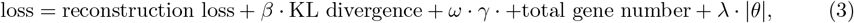

where we used 4 weights, *ω* = 1, 1.5, 2, 2.5 to test the loss function. We also explored using the same loss function but with a different gamma that is linearly annealed from 2.0 to 0.1. L1 regularisation was also used.

#### 4.1.5 Training

The data were split into training, validation and test sizes of 4167, 1190, 596 respectively. A basic hyperparameter optimisation was performed on model v0 using a grid search over three parameters:

- Hidden Dimension: tested with values of 256, 512, and 1,024.
- Latent Dimension: tested with values of 32, 64, and 128.
- Learning Rate: tested with values of 0.01 and 0.001.

All models used the early stopping regularisation technique. This technique prevents the model from overfitting by monitoring its performance on a validation set and halting the training process if the performance does not improve to preserve its generalisation capability. We also explored dropout but found it did not improve performance in this scenario. The epoch number was set at 10,000 for all models. The final model architecture is shown in Figure S6.

The model training was assessed by measuring training and validation losses over the trained epochs. F1 score was calculated to assess the model’s performance in the test data. The latent space of the VAE models was used to validate and explore the results, especially in the basic VAE model. Such analysis allowed us to verify the model’s ability to explore latent space and see valuable patterns that the VAE encoded in the data of interest.

### 4.2 Sampling

To generate new genomic sequences, we sampled 10,000 latent vectors from a multivariate normal distribution whose dimensionality matched the model’s latent space. We assumed a standard normal distribution with a mean vector of zero and an identity covariance matrix. Each sampled latent vector was then passed through the decoder of the corresponding model to generate new genomic sequences. Such an approach allowed us to assess the diversity of the genomes that the model can produce.

### *4*.*3 E. coli* whole-cell model simulations

To evaluate the viability and physiological consequences of our minimal-genome designs, we used a single-cell *E. coli* whole-cell model (WCM) as an *in silico* validation step [30]. In brief, the WCM can be viewed as a large system of coupled ordinary differential equations in which cellular states (e.g. metabolite pools, macromolecule counts, chromosome configuration) are updated by process-specific dynamical rules. Two key processes, RNA/protein synthesis and degradation, are treated stochastically using multinomial and Poisson sampling, respectively, while shared resources are partitioned between processes at each timestep to enforce mass conservation. The model is implemented in Python with Cython-optimised inner loops and is available at https://github.com/CovertLab/wcEcoli, and for this work we used the version which was available as of February 2023. The model integrates data from three distinct lab strains (K-12 MG1655, B/r and BW25113) and therefore represents a composite “hybrid” *E. coli* strain with 1,872 modelled genes.

Simulations were configured to follow the default log-phase growth condition in the WCM, representing M9 minimal aerobic medium supplemented with 0.4% glucose at 37. Under these conditions the model cells grow and divide for multiple generations, where one generation denotes a complete cell-division cycle. To balance runtime with memory and storage constraints while still allowing steady-state behaviour to emerge, we limited simulations to a maximum of 20 generations and defined a strain as *WCM-viable* if the corresponding simulation completed all 20 generations without numerical failure. The WCM uses the FireWorks workflow management tool to manage and dispatch simulation jobs, allowing us to execute all simulations in parallel on the University of Bristol’s BlueCrystal phase 4 supercomputer. Unless otherwise stated in the Results or Supplementary Information, simulations were run in triplicate with independent random seeds.

For the purposes of this study we defined the “WCM gene set” as the 1,872 genes explicitly modelled in the WCM. Each genome considered in this work (lineage representatives and VAE-sampled genomes) is represented as a binary presence–absence vector over the pangenome, and prior to *in silico* validation, we restricted this vector to the WCM gene set. Since the WCM requires gene deletions to be specified explicitly, we then identified, for each genome, the subset of WCM genes absent from a given vector and implemented these as simultaneous knockouts in the model. A knockout of gene is implemented by forcing its final transcriptional expression to zero, which in turn sets its transcription initiation probability and regulatory baselines to zero, removes the corresponding mRNA species and prevents production of the encoded protein(s).

To calibrate and interpret WCM-predicted essentiality, we first performed an *in silico* genome-wide single-gene knockout screen. For this, each gene in the WCM gene set was deleted individually under the same environmental conditions and 20-generation simulation window as above, following which, a gene was classified as *WCM-essential* if its deletion prevented the simulated cell from completing all 20 generations. We compared these classifications to CRISPR-knockout-based fitness measurements from Rousset *et al*. [31], yielding 90.2% accuracy (95.3% specificity, 51.6% sensitivity). This provides experimental grounding for using the WCM as a computational screening tool for minimal genome designs, while acknowledging that it does not validate higher-order synergistic effects of combinatorial deletions.

For each lineage or VAE-sampled genome used in downstream analyses, we then applied the following procedure: first, restrict the genome’s presence–absence vector to the WCM gene set, treating all absent WCM genes as candidate deletions, then we restored any genes belonging to the WCM-essential set, enforcing their presence irrespective of the original genome vector, and finally, we applied the resulting gene knockout pattern to the WCM and simulated growth under the standard protocol described above. This ensures that non-viability arises from the interaction of non-essential deletions within the mechanistic WCM rather than from the trivial absence of genes that are essential by construction of the model.

### 4.4 Genome realization algorithm

An algorithm was developed to generate new minimised genomes that loads a wild-type sequence of the K12 MG1655 *E. coli* genome and a list of genes that must be present in each final genome. The algorithm then extracts non-essential genes to be removed from the wild-type sequence by comparing whether the gene is present in the list of genes that must be present in the sequence of interest. When non-essential gene features are extracted, their corresponding nucleotide positions are marked for removal. However, all the essential genes are still kept in the sequences. The algorithm then generates the minimised genomes by removing the positions marked for removal and removing genetic segments while keeping the other features intact. The algorithm saves the generated sequence in a Fasta file.

## 5 Data and code availability

The code can be accessed from: https://github.com/ucl-cssb/genome-minimizer-2.

Data used or generated during this project can be accessed from Zenodo: 10.5281/zenodo.17760396

