## Supplementary Information for "Designing minimal *E. coli* genomes using variational autoencoders"

### Designing minimal *E. coli* genomes using variational autoencoders: Supplementary information

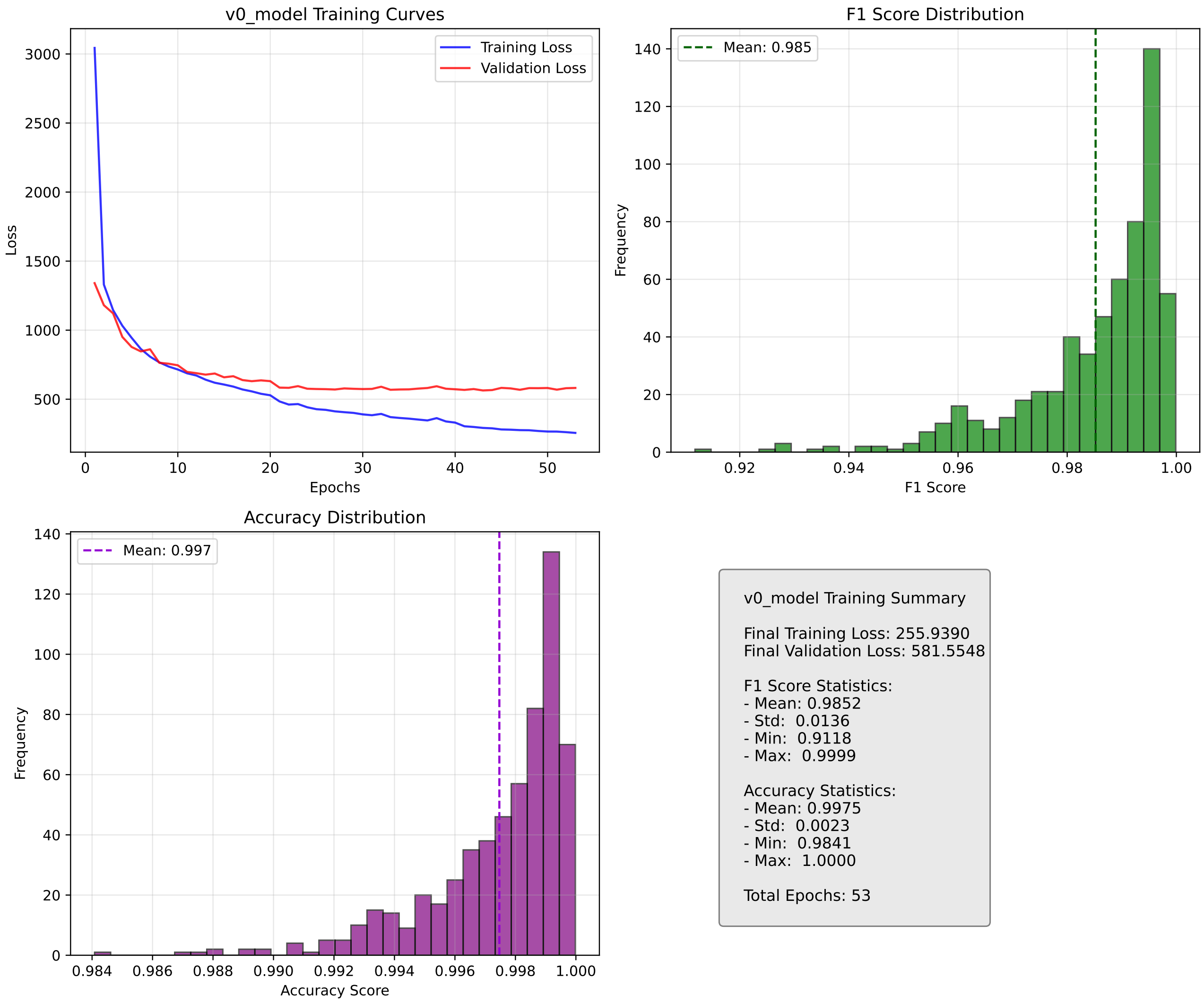


Figure 1: Model training summary for model version v0

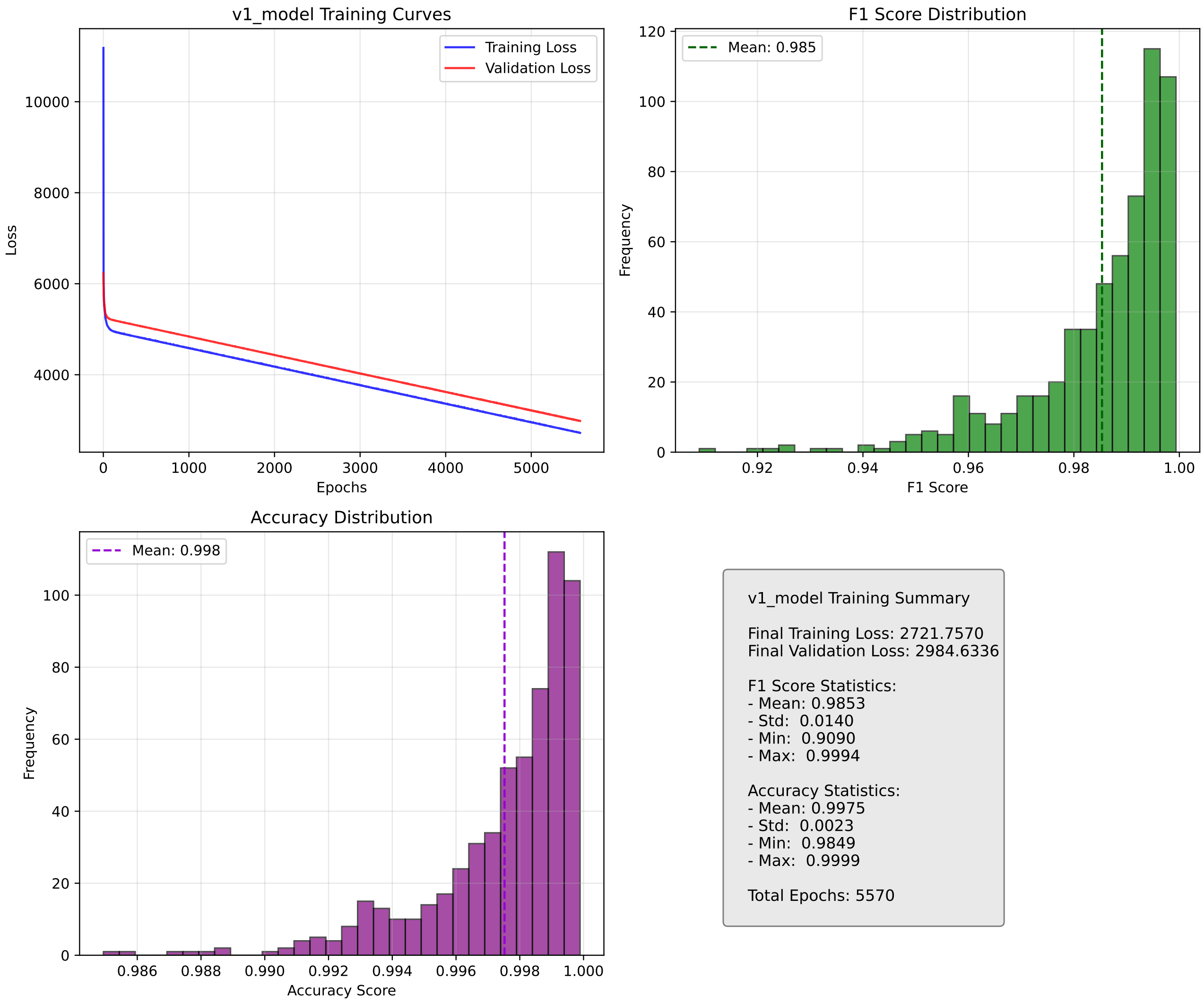


Figure 2: Model training summary for model version v1

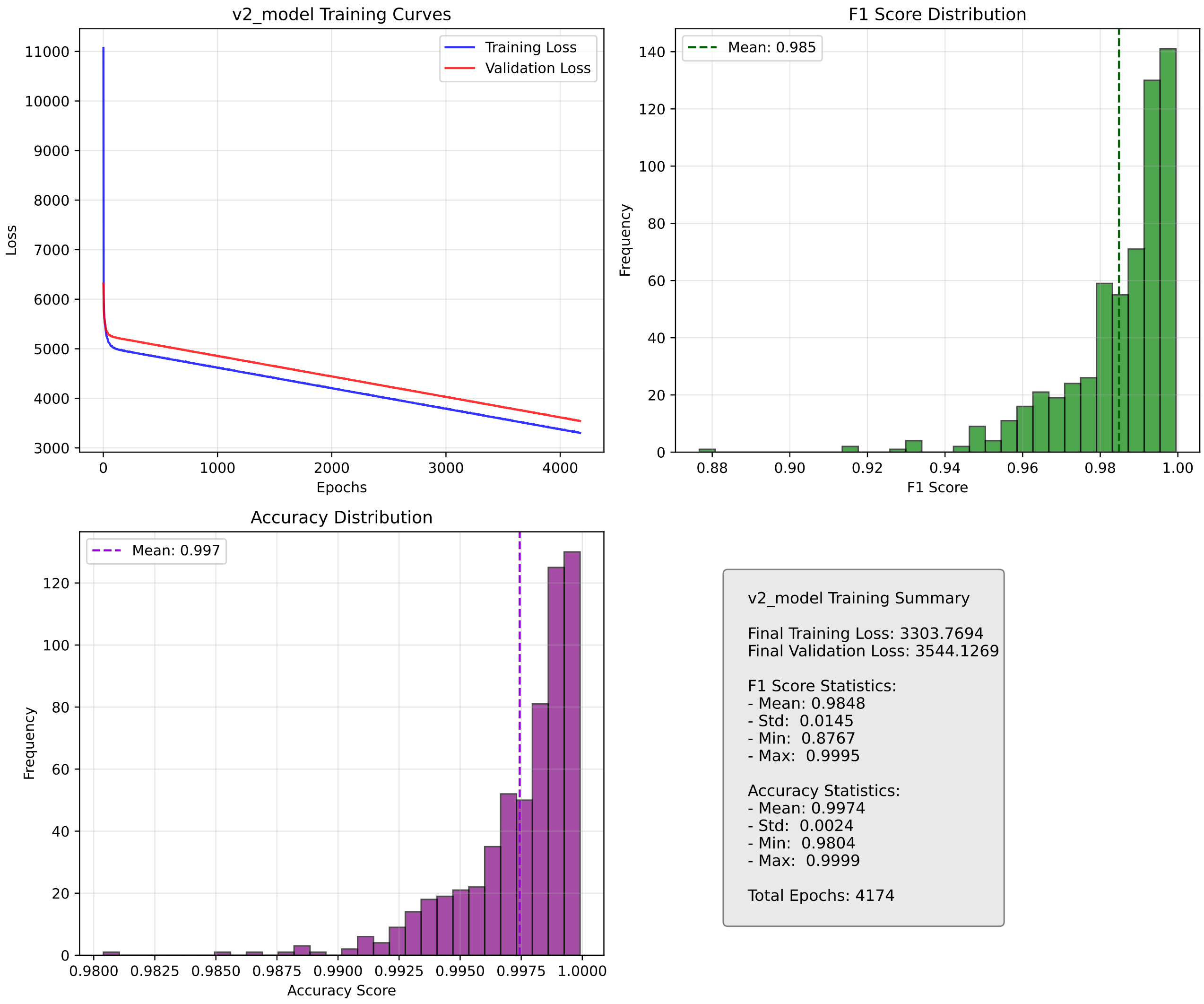


Figure 3: Model training summary for model version v2

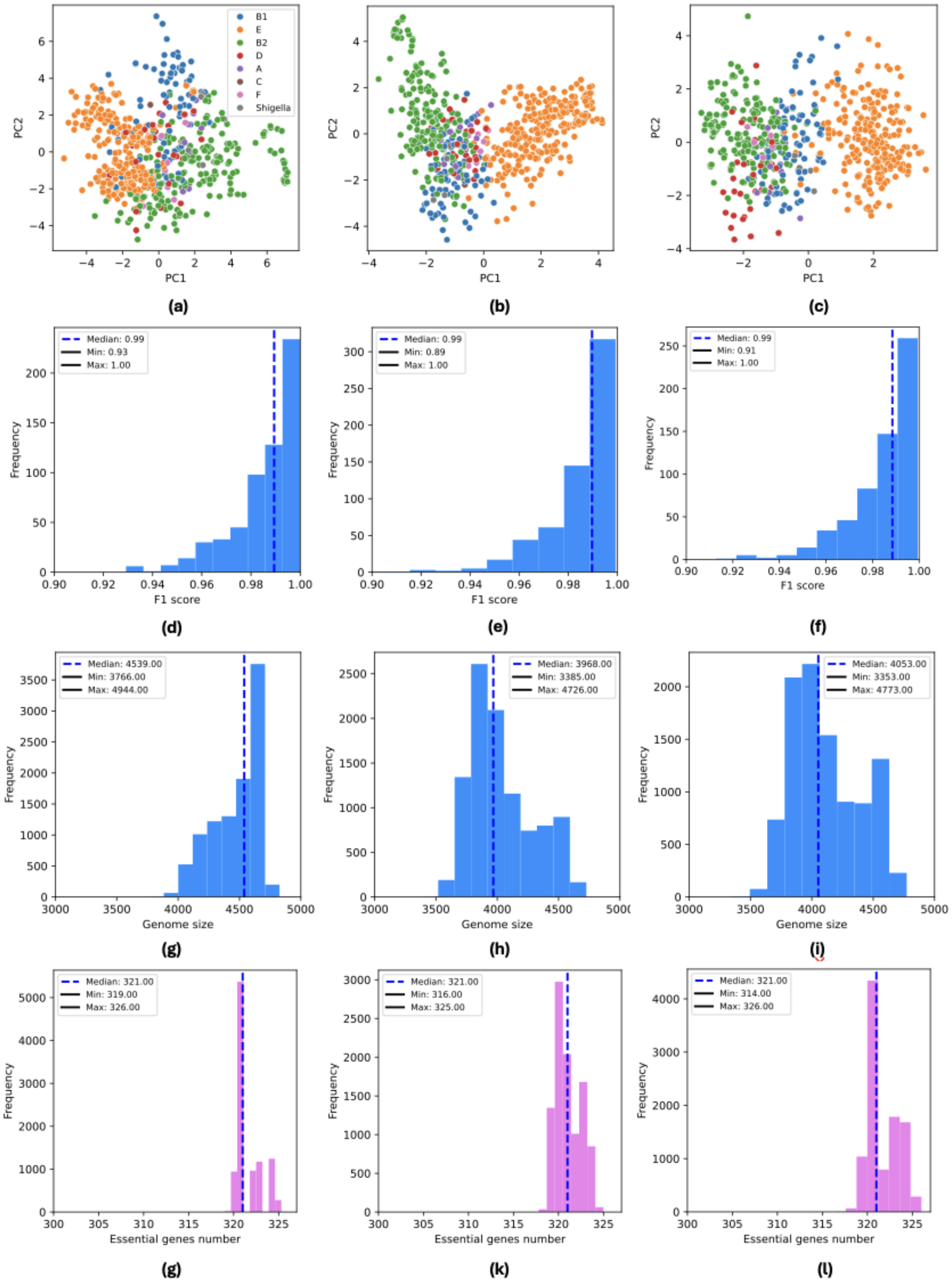


Figure 4: Model development. Latent space and distributions of F1 score, genome size and essential gene number for models of increasing complexity; v0 (a, d, g, j), v1 (b, e, h, k) and v2 (c, f, i, l).

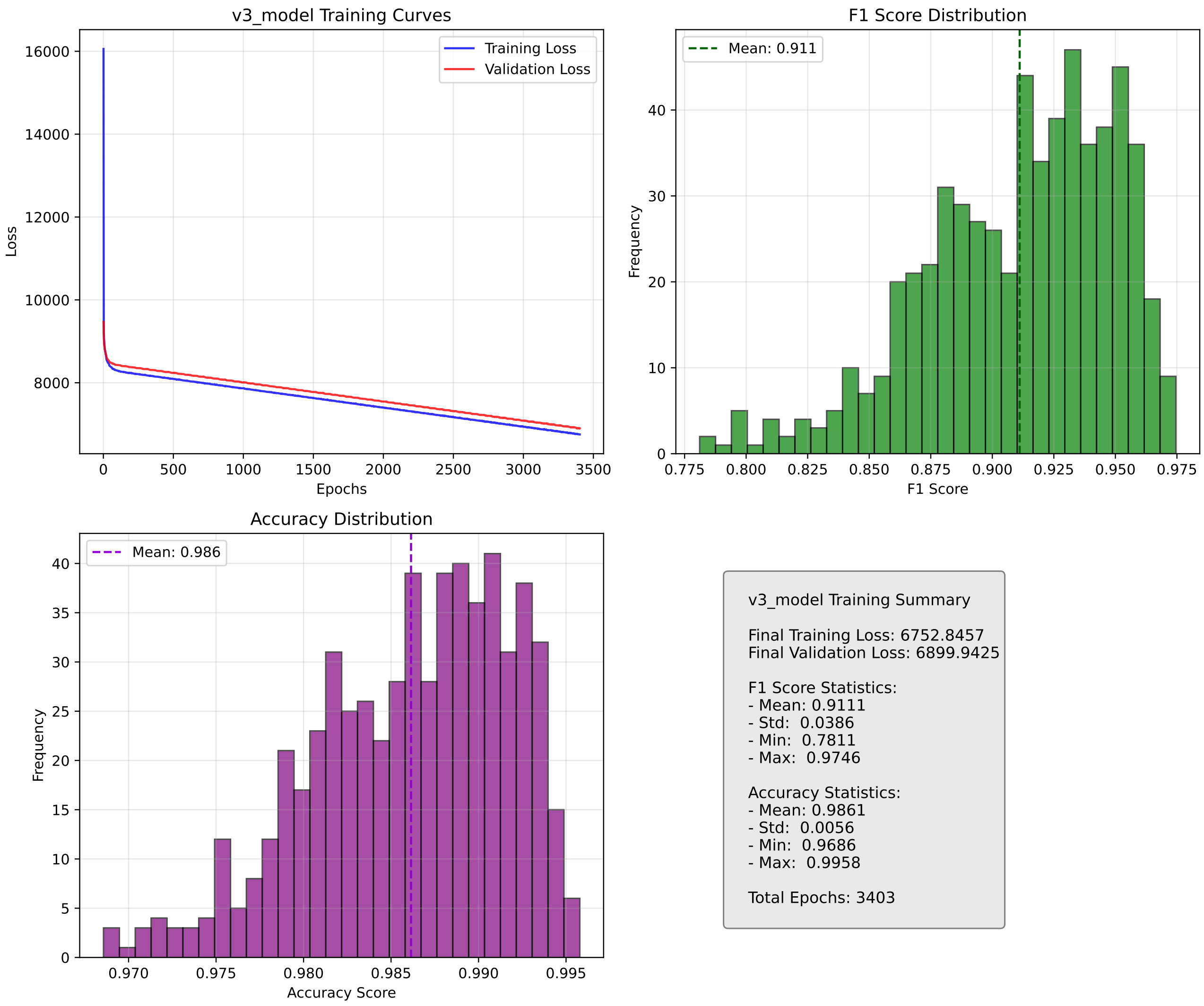


Figure 5: Model training summary for model version v3

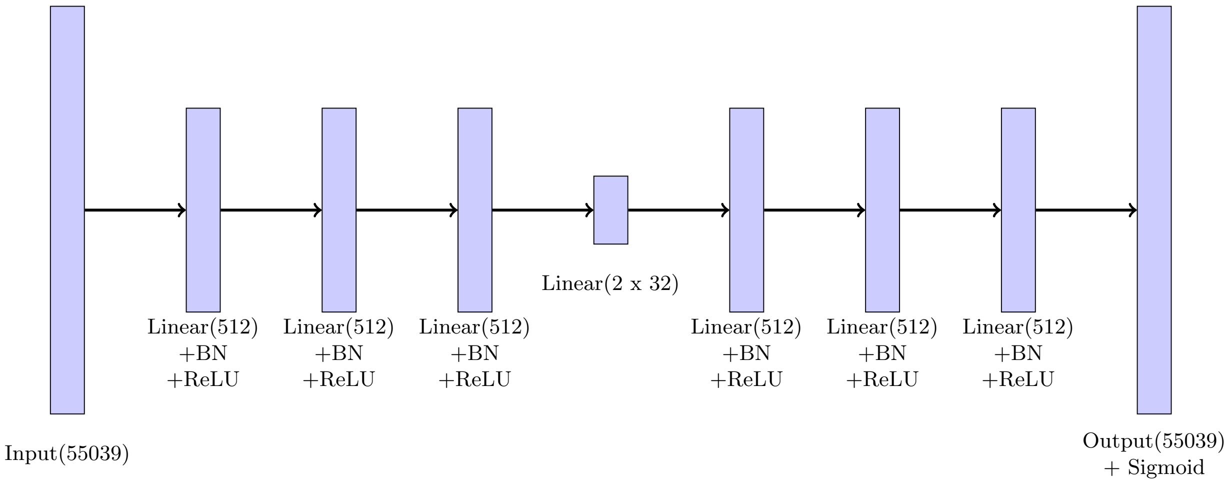


Figure 6: Implemented VAE architecture.
